# Identifying differential effects from eleven mixing techniques on mRNA lipid nanoparticle physicochemistry and biological performance

**DOI:** 10.1101/2025.11.07.687311

**Authors:** T. Bethiana, A. Aljabbari, Y. Li, H. Mitra, M. Baghbanbashi, G. Harris, S.R. Dasaro, F. Masoomi, F. S. Vago, S. L. Hartzler, M. Figueiredo, L. A. Metskas, P. Vlachos, A. Ardekani, Y. Yeo, K. Ristroph

## Abstract

Lipid nanoparticle (LNP) formulation requires a mixing step. Many studies, especially from academic groups, utilize either microfluidic mixers or hand mixing to prepare LNPs, but commercial-scale processes use turbulent-flow mixers. This discrepancy in mixing techniques has been underexplored, as LNPs made by different techniques may exhibit different performance, such that bench-scale results cannot be replicated using materials manufactured at scale. We here isolate and interrogate the effect of primary mixing. Lipid nanoparticles are produced from ten mixers (one used in two ways), holding all other formulation parameters constant, to directly compare across techniques. LNPs produced from the different mixers exhibit widely different physical properties and biological performance. Notably, manual pipetting common in academic practice yields particles that do not resemble those produces by turbulent-flow mixers. Findings are connected mechanistically to physicochemical characteristics that arise from the different flow regimes. Further establishing the relationship between mixing and LNP properties is critical.

**Graphical abstract:** 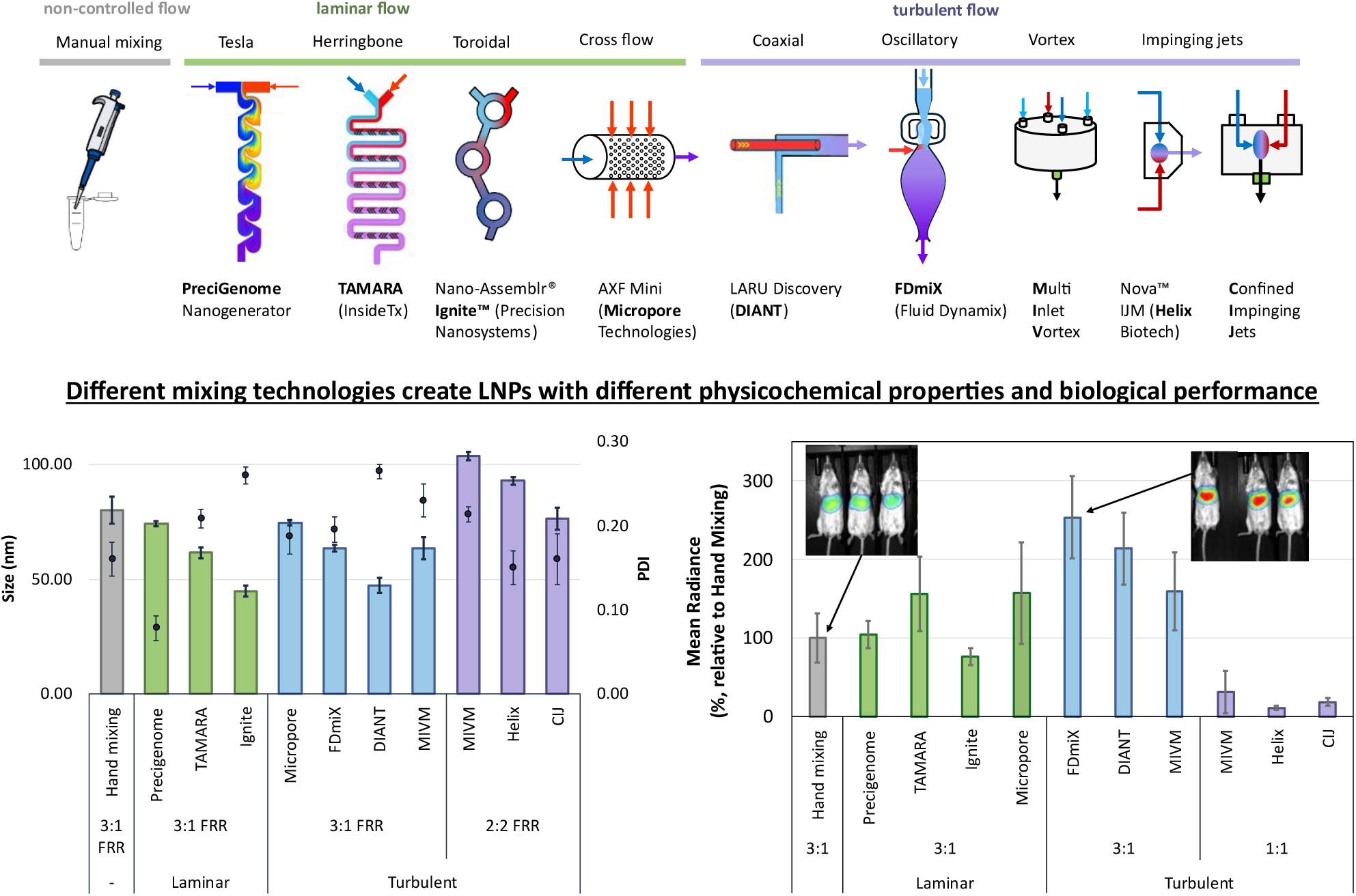

## Body

Lipid nanoparticles (LNPs) have rapidly become the non-viral platform of choice for nucleic acid delivery.^1–3,4–6^ Significant challenges remain in the reproducibility and translation of LNP formulations from laboratory-scale discovery to clinical manufacturing.^7,8^ A key difference between these scales is the mixing technique. The most common formulation techniques at the bench scale use either microfluidic or hand mixing to prepare LNPs, but at commercial scale – e.g., for the COMIRNATY® vaccine – turbulent-flow mixers are used.^9^ A deeper understanding of the LNP self-assembly process as a function of flow and mixing during formulation, which is often overlooked in formulation optimization, is needed to bridge this gap.^10–13^

RNA LNP self-assembly occurs when an ethanol solution containing dissolved lipids is rapidly mixed with an aqueous buffer containing nucleic acid. The solvent exchange drastically lowers lipid solubility, leading to supersaturation that drives precipitation. As this process is kinetically controlled, the timing of solvent dilution, ionizable lipid–nucleic acid complexation, and particle nucleation is highly sensitive to the speed of mixing, which in turn is a function both of mixer geometry aregime.^14^ Mixing that is sufficiently faster than nanoparticle self-assembly is required to produce homogeneous particles.^9^ LNP self-assembly is estimated to occur over tens of milliseconds; techniques that take longer than this to mix water and ethanol, such as the manual pipetting and microfluidic mixing frequently used at bench scale, therefore may not produce the same LNPs that would result from the same inlet streams being mixed by turbulent mixers at commercial scale.^9^

In this paper, eleven mixing techniques, including turbulent, microfluidic, and manual mixing are evaluated with other variables held constant such that all formulations are compositionally identical. The LNPs produced exhibit widely different physical properties – particle size distribution, encapsulation efficiency, and internal structure – and biological performance as a function of mixing. In particular, LNPs made by turbulent mixing outperform the most common bench-scale microfluidic and manual mixing techniques. For researchers working in the LNP formulation optimization space – e.g. designing new ionizable lipids, incorporating biodegradable linkers, and optimizing PEG densities – these results suggest that continued reliance on non-scalable mixing will introduce major experimental artifacts into the structure and performance of the resulting lipid nanoparticles that will hinder translatability. Aligning bench-scale practice with clinically relevant manufacturing processes is therefore critical to ensure that findings at small scale can be reproduced in large-scale commercial manufacture.

### Mixers, scales, and flow regimes

Mixing technologies for LNP production that incorporate different mixing flow conditions have diversified substantially in recent years. Early benchtop studies relied on mixing by manual pipetting, a convenient but loosely controlled technique. The limitations of hand mixing motivated the commercialization of the toroidal microfluidic mixing platform NanoAssemblr Ignite, which produces standardized laminar mixing conditions and has been widely adopted for laboratory use.^15–17^

A pair of turbulent mixers, the Confined Impinging Jets (CIJ) mixer and Multi-Inlet Vortex Mixer (MIVM), were developed and studied for polymeric and lipid nanoparticle formulation for the past two decades.^18–20^ Impinging jets mixers are used in the global-scale manufacture of COMIRNATY®, the Pfizer-BioNTech vaccine against SARS-CoV-2, and Moderna’s patent portfolio includes processes that utilize the MIVM.^9^ The CIJ and MIVM operate at high Reynolds numbers, which produces much faster homogenization times than laminar mixing and ensures mixer scalability.^9,21^ A drawback of these operating conditions is the relatively high minimum material requirement for lab-scale work compared to microfluidics.

Several new mixers, designed to work either in laminar or turbulent flow, have recently entered the market. DIANT introduced a continuous co-axial jet system shortly after its founding in 2019.^8^ FDX Fluid Dynamix developed a mixer employing oscillatory jet technology, the FDmiX, which was incorporated into Lonza’s nucleic-acid encapsulation GMP workflows in 2024. Helix Biotech released a modular Nova™ impinging jet mixer.^22^ Other laminar-flow mixers include PreciGenome’s NanoGenerator, which uses a tesla microstructure, and Inside Therapeutics’s TAMARA mixer which uses staggered herringbone laminar geometries.

Laminar systems that rely on diffusion-limited mixing are unsuitable for large-scale manufacturing because of fouling and the requirement for parallelization to scale out rather than true scale-up.^23–25^ They remain favored in academic settings due to their affordability, operational simplicity, and rapid turnaround for exploratory studies on lipid chemistries and formulation optimization.^3,8,26^ Industrial pipelines have focused on turbulent systems for greater scalability and robustness.^7^ Recent studies have highlighted mixing as one of the most sensitive yet most variable steps in LNP production, with most investigations to date concentrated on parameter optimization within individual devices or on scaling microfluidic approaches.^26,27^ To the authors’ best knowledge, there has been no comprehensive head-to-head comparison across the broad spectrum of commercialized laminar and turbulent mixers.

### Mixer operating conditions

Eleven mixing techniques were compared: hand mixing, and nine commercially available mixers (one used in two modes). Mixers were operated under flow conditions recommended by their manufacturers. The same ethanol feed stream composition was used in all eleven cases. The eight conditions tested at a 3:1 buffer:ethanol FRR all had identical solids concentrations, pH and ionic strength, and solvent quality at the point of mixing. The three conditions tested at a 1:1 FRR also had matching solids concentrations, pH and ionic strength, and solvent quality at the point of mixing. All eleven conditions had identical solids concentrations, ionic strength, solvent quality, N/P ratio, and pH following the downstream quench step, which was performed 5 seconds after collection was completed to avoid introducing artifacts from quenching LNPs prematurely before particle formation at the conditions inside the mixer was complete.

LNPs produced using hand mixing served as the benchmark condition for evaluating other mixers. To assess internal reproducibility across the mixing technologies and test conditions, LNPs with yeast RNA as the payload were first produced and characterized (size, PDI, zeta potential, and EE). Once reproducibility was established, LNPs encapsulating FLuc mRNA (mRNA encoding firefly luciferase) were produced and evaluated using a comprehensive suite of tests expanded to include internal structural analysis and biological performance (workflows, **Figure S1**).

Mixers operated with syringe pumps allocated a small pre-waste volume to ensure that flow reached steady state before samples were collected. A post-waste was also discarded in some cases to avoid collecting fractions that had aged differently than the main sample. By excluding both pre-waste and post-waste fractions, the collected samples represent nanoparticles generated under reproducible mixing conditions, rather than artifacts arising from startup effects or time-dependent evolution.

### Yeast RNA LNP production and characterization

Seven independent replicates (using new stock solutions each time) of LNPs loaded with yeast RNA were prepared using each mixing technology to determine replicability. Consistently narrow error bars in the results of the physicochemical characterization were observed (**Figure 1**), indicating that each mixer can deliver stable and controllable mixing conditions. Variability observed in particle size, surface charge, and encapsulation efficiency across samples is therefore primarily driven by mixing geometry or FRR, rather than inconsistencies in the individually applied mixing processes. Statistical differences between the physicochemical characteristics of LNPs made by each mixer were assessed using Tukey’s HSD test and pairwise comparisons of all parameters. This analysis led to ten clusters visualized by compact letter displays as “A”, “AB”, “B”, “BC”, “C”, “CD”, “D”, “DE”, “E”, and “F” as shown in **Figure 1a-d**. Any two treatments receiving the same letter at the top of the graphs are not significantly different as per Tukey’s HSD method.

**Figure 1.**
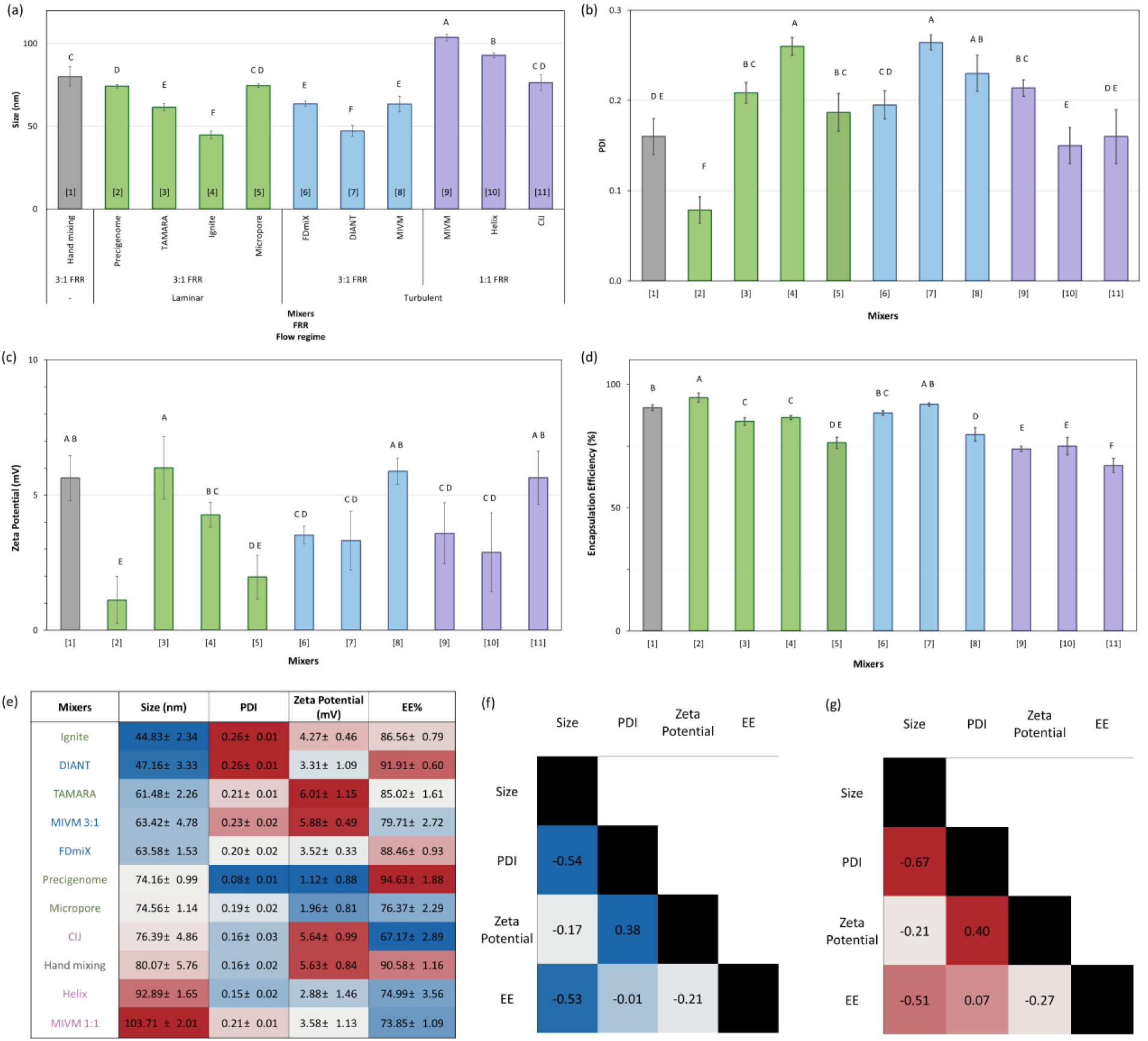
The Compact Letter Display (CLD) graphs portray pairwise differences between (a) z-average diameter, (b) polydispersity index, (c) zeta potential, and (d) encapsulation efficiency. In each CLD graph, 7 data points were averaged and presented along with respective standard deviations. (e) Statistical analysis of yeast RNA LNP physicochemical characteristics (size, PDI, zeta potential, and encapsulation efficiency) visualized as a heatmap, sorted by particle size in ascending order. Blue indicates low values, white indicates median values, and red indicates high values. Each box represents an average of seven data points; (f) Pearson correlation matrix, where blue shading denotes stronger linear associations and white denotes weaker associations; (g) Spearman correlation matrix, where red shading denotes stronger monotonic relationships and white denotes little to no monotonic association. The mixer names are color coded: grey text denotes an undefined flow regime at 3:1 FRR, green text denotes laminar flow mixers at 3:1 FRR, blue text denotes turbulent flow mixers at 3:1 FRR, and purple text denotes turbulent flow mixers at 1:1 FRR.

The LNPs produced at a 1:1 flow rate ratio produced the largest LNPs, [average] vs [average] produced at a 3:1 FRR across all mixers. The observed effect of solvent quality on particle formation is also captured by the operation of the same mixer, the MIVM, at FRR 3:1 versus 1:1 (size vs size). The effect of solvent quality was studied in this way because the impinging jets mixer used commercially is designed to operate at a 1:1 FRR, despite the popularity of a 3:1 FRR in the literature.^28,29^ The most common lab-scale operation of the CIJ includes an immediate downstream quench by collecting effluent directly in a vial containing buffer.^48^ When this is done, LNPs formed at a 1:1 FRR are much smaller than what was observed here.^30^ This suggests the importance of an immediate downstream quench when formulating LNPs at a 1:1 FRR to trap a kinetic state during formation, corroborating a recent report.^28^ Standardizing and optimizing the mixing for this quench step is an area for further development.

Among the eight mixers tested at a 3:1 FRR with identical compositions at the point of mixing, significant differences were observed in LNP size, PDI, zeta potential, and encapsulation efficiency. The microfluidic Ignite and turbulent DIANT mixers produced the smallest LNPs but with the largest PDIs. PDIs from all LNPs were below 0.3, which is acceptable for drug delivery applications.^31^ Although all LNPs exhibited neutral zeta potential values between –10 and +10 mV, statistically significant differences in zeta potential were observed, suggesting subtle differences in lipid organization and surface presentation.

Mixers operated at a 3:1 FRR generally achieved higher encapsulation efficiency than those operated at 1:1. Of the eight mixing conditions, Precigenome and DIANT produced the highest EEs, exceeding 90%, while EEs produced at 1:1 FRR were closer to 70%. The impact of aqueous-to-organic ratio results on their encapsulation efficiency is likely due to more favorable lipid–RNA assembly at higher dilution conditions, consistent with previous studies.^5^

The relationships among physicochemical characteristics of yeast RNA LNPs produced by different mixers are summarized as a heatmap in **Figure 1e**; by Pearson correlations (r) (**Figure 1f**) in which values closer to ±1 indicate stronger associations; and by Spearman correlations (ρ) (**Figure 1g**) in with values closer to ±1 reflect strong monotonic relationships and values near 0 indicate no monotonic association. An inverse correlation between particle size and encapsulation efficiency was observed in which smaller particles consistently exhibited higher RNA encapsulation (r = –0.53, ρ = –0.51). Size and PDI also appeared inversely correlated, with larger LNP populations exhibiting lower PDI and smaller particles a higher PDI (r = –0.54, ρ = –0.67), consistent with previous findings.^6^ PDI and zeta potential showed a moderate correlation (r = +0.38, ρ = +0.40), with low-PDI LNPs from mixers such as TAMARA exhibiting higher zeta potential and higher-PDI LNPs from Ignite, DIANT, and CIJ exhibiting lower values.

### FLuc mRNA LNP production and characterization

We then replicated the operational conditions and suite of mixing technologies discussed above to produce LNPs encapsulating FLuc mRNA to enable evaluation of transfection efficiency (production workflow **Figure S1**). After production, LNPs were dialyzed against HEPES buffer to remove ethanol and adjust the pH to 7.4. LNP sizes shift during this step because of particle fusion due to the loss of electrostatic colloidal stability above the pKa of the ionizable lipid.^22,32,33,34,6,35^ Dialyzed LNPs were concentrated by centrifugal ultrafiltration prior to evaluation by SAXS, cryo-TEM, the TNS assay, and protein expression *in vivo*. Physicochemical characterization results are presented as heatmaps in **Figure 2, Figure 3**, and **Figure 4** for samples after formulation (‘pre-dialysis’), post-dialysis, and post-concentration, respectively.

**Figure 2.**
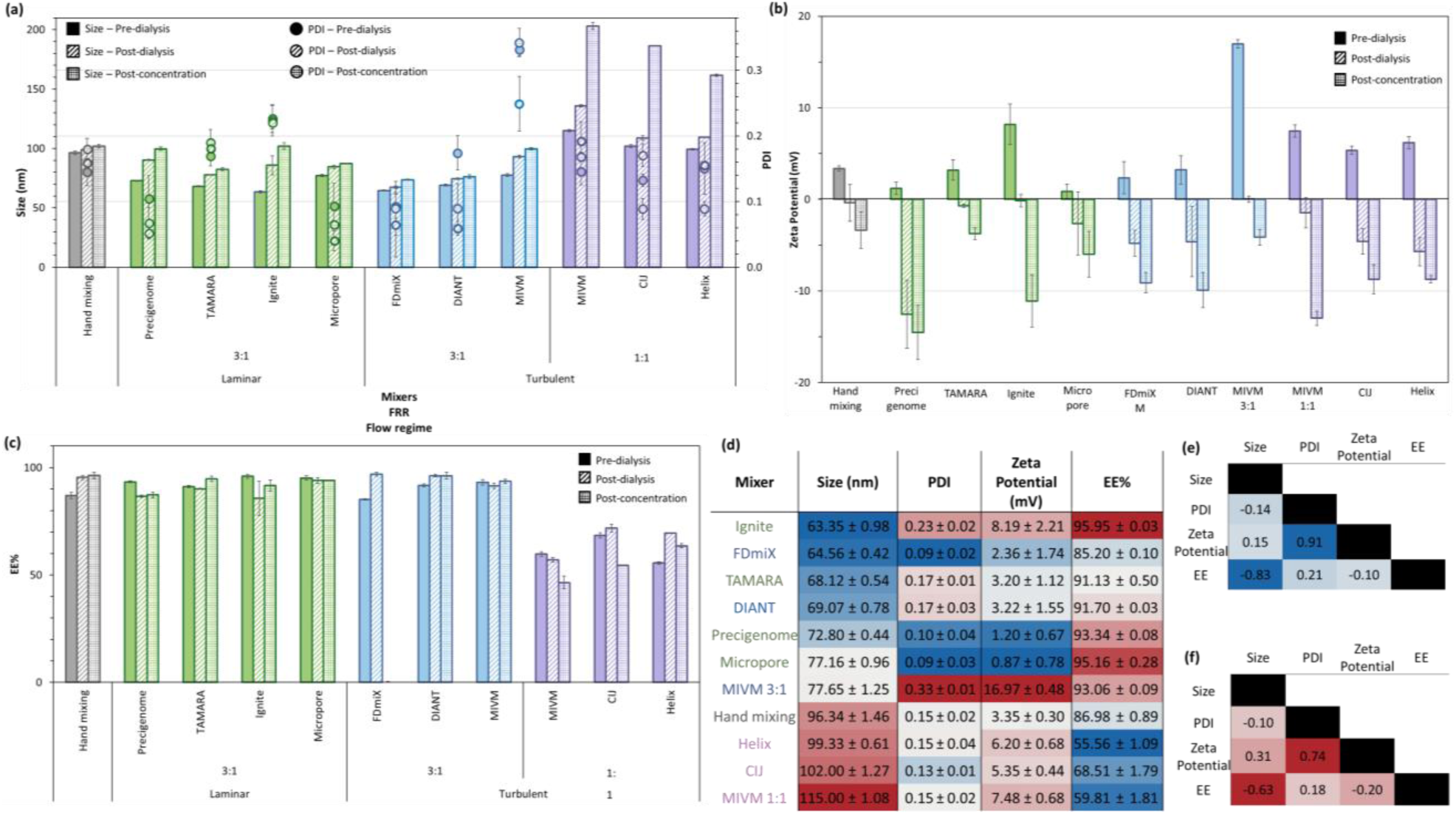
(a) Size (bars) and PDI (dots) measurements on pre-dialysis LNPs; (b) Zeta potential measurement on pre-dialysis LNP; (c) Encapsulation efficiency by Ribogreen assay measurements on pre-dialysis LNPs; (d) Heatmap analysis for physicochemical properties of pre-dialysis FLuc mRNA LNPs sorted by size. Standard deviations reflect variation from statistical replicates.

**Figure 3.**
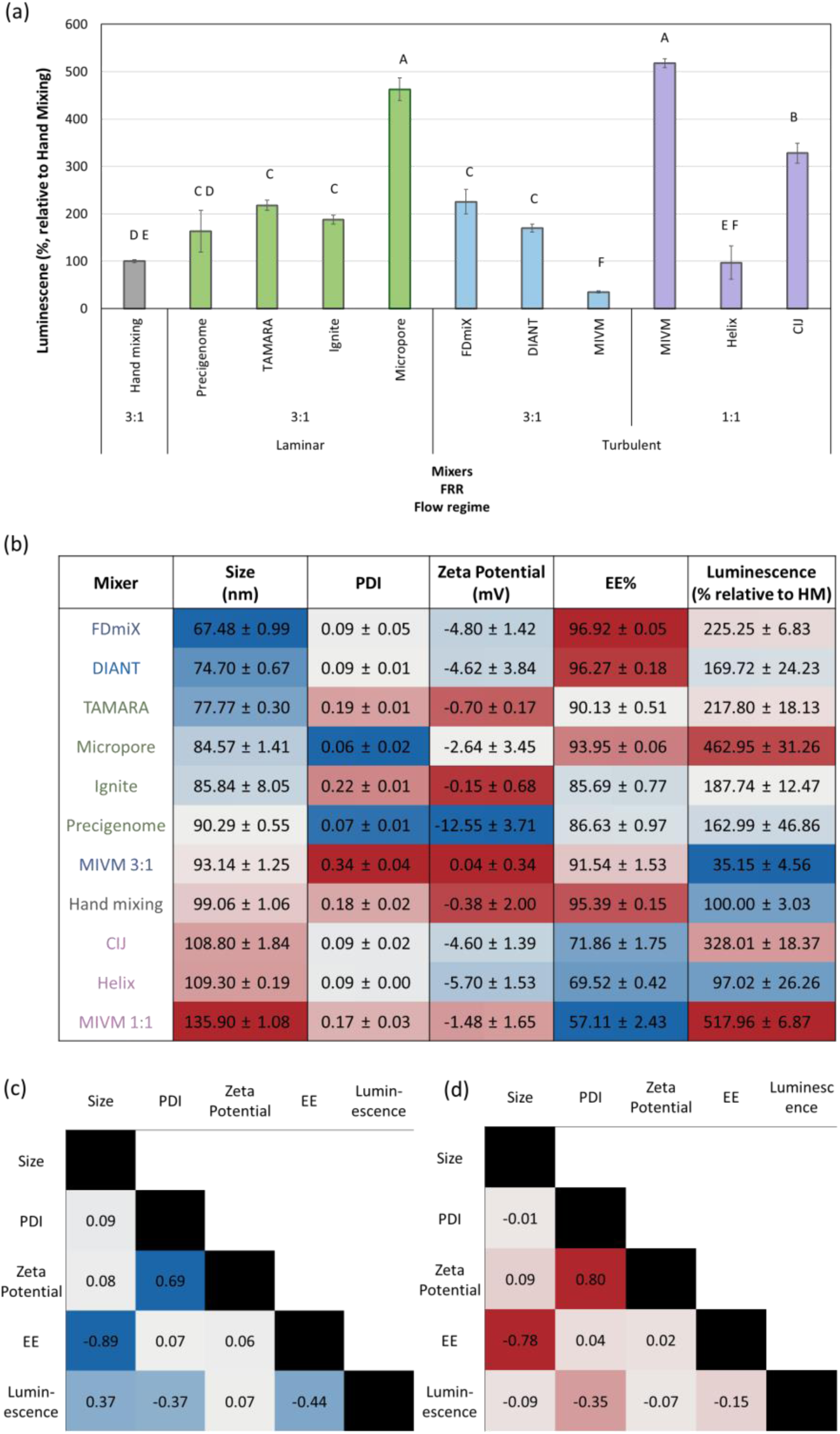
(a) Luminescence of FLuc mRNA LNPs prepared with different mixers, normalized to hand mixing; (b) Heatmap comparison of post-dialyzed FLuc mRNA LNP properties relevant to *in vitro* test across different mixers, sorted by size. (c) Pearson correlation matrix, where blue shading denotes stronger linear associations and white denotes weaker associations. (d) Spearman correlation matrix, where red shading denotes stronger monotonic relationships and white denotes little to no monotonic association. Standard deviations reflect variation from statistical replicates.

**Figure 4.**
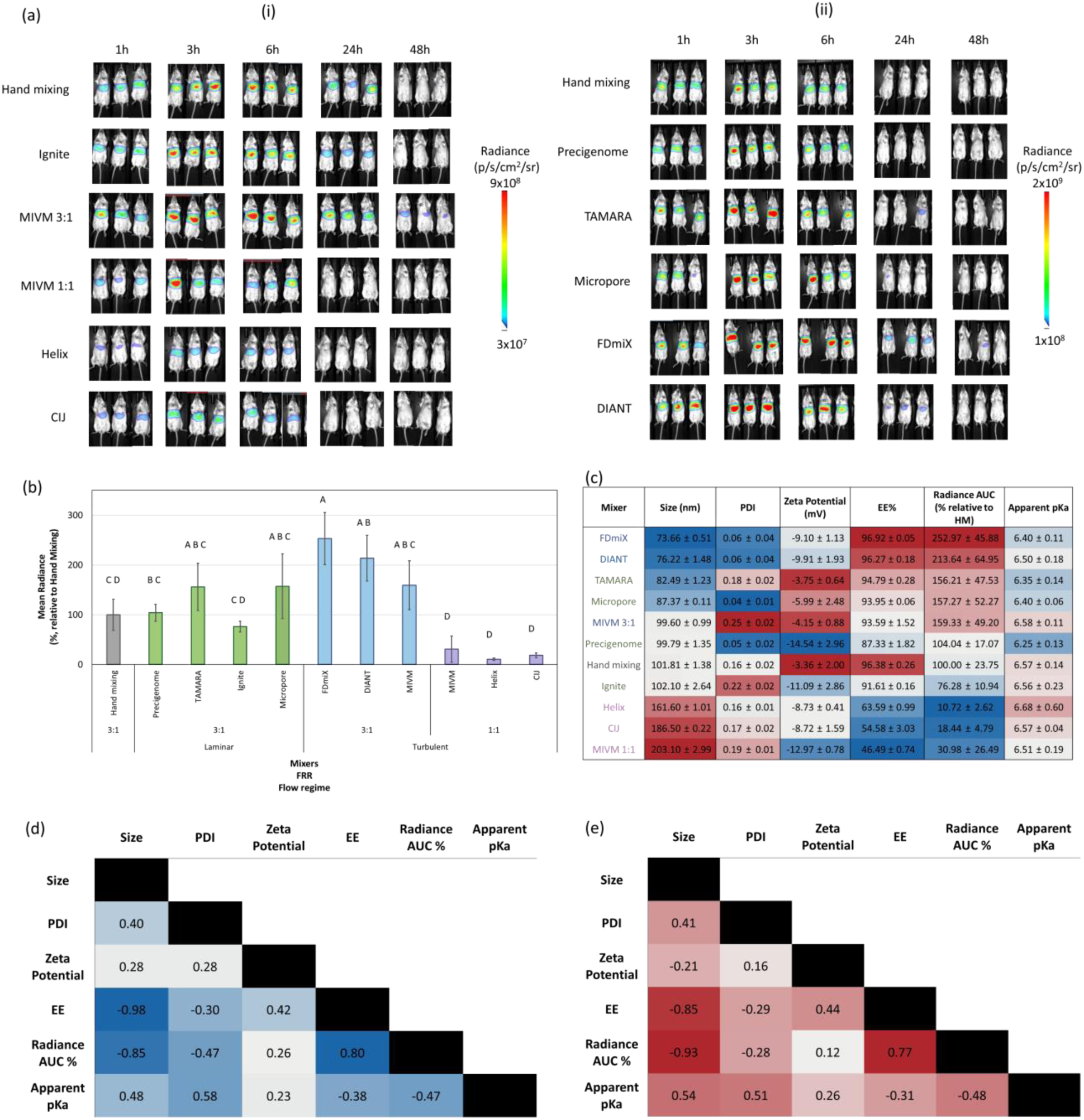
(a) In vivo bioluminescence imaging of FLuc mRNA LNPs formulated with different mixers. 7 week-old Balb/c mice (n=3/group) received LNPs at 0.2 mg/kg mRNA by IV injection. The mice were imaged with IP injected luciferin (50 mg/kg) at 1, 3, 6, 24, and 48 h after the LNP injection. Luminescence intensity (p/s/cm²/sr) is shown on a pseudo color scale (blue–red), indicating relative luciferase expression. (i) First round of 6 mixers, (ii) Second round of 6 mixers; (b) Comparison of mean radiance (x 10^8 photons/s/cm²/sr) area under the curve (AUC) averaged over 5 timepoints with CLD; (c) Heatmap comparison of post-concentrated LNP properties across mixers, including parameters relevant to in vivo delivery and apparent pKa. (d) Pearson correlation matrix, where blue shading denotes stronger linear associations and white denotes weaker associations. (e) Spearman correlation matrix, where red shading denotes stronger monotonic relationships and white denotes little to no monotonic association. Standard deviations reflect variation from statistical replicates.

LNPs produced using FLuc mRNA exhibited physiochemical properties and trends that mirrored those produced using yeast RNA. Notably, the pattern of smaller LNPs at FRR 3:1 than FRR 1:1 persisted and the inverse correlation between size and encapsulation efficiency was observed (r = –0.83, ρ = –0.63). The LNPs produced at a 3:1 FRR via any mixer device were smaller than those produced at a 3:1 FRR via hand mixing (63 – 77 nm, vs 96 nm). PDI was more strongly positively correlated with zeta potential than in the case with yeast RNA LNPs (r = 0.91, ρ = 0.74), and particle size was no longer correlated with PDI (r= –0.14, ρ = –0.10).

### *In vitro* mRNA translation

HEK293 cells were treated with FLuc mRNA-loaded LNPs at 0.65 μg/mL of mRNA in the medium. Total mRNA concentration was determined using the Ribogreen assay and applied as the basis for both *in vitro* and *in vivo* studies. Dosage definition by total mRNA concentration aligns with preclinical and clinical studies, which report mRNA doses without correcting for encapsulation efficiency.^36^ In well-optimized formulations, encapsulation efficiency is typically close to 100%; therefore, total mRNA concentration approximates encapsulated mRNA concentration. As a result, EE is generally treated as a formulation quality control attribute, while dosing in preclinical and clinical studies is defined by total mRNA administered.^37^ Encapsulation efficiency did not directly correlate with *in vitro* protein expression, as some formulations with lower EE still produced strong luminescence at the same total mRNA dose (Figure 3). This demonstrates that while EE is commonly used as a LNP formulation quality control attribute, it does not necessarily predict functional performance *in vitro*. Instead, the *in vitro* biological activity reflects the combined influence of physiochemical properties of LNPs and their subsequent intracellular fate. Significant differences were observed in the level of protein expression induced by the LNPs made by the different mixers. LNPs formulated using the Micropore (3:1, laminar) and MIVM 1:1 (turbulent) demonstrated the strongest in vitro signals, producing over 5-fold higher expression relative to hand mixing (Figure 3a). All other laminar mixers showed moderately higher expression than hand mixing and similar performance to the FDmiX and DIANT.

### *In vivo* mRNA translation in relation to apparent pKa of LNPs and endosomal escape

Products from all 11 mixing techniques could not be tested in a single experiment, so the in vivo study was split into two rounds, each testing LNPs produced using six techniques, with hand mixing as the common benchmark. Post-dialysis, post-concentration LNPs (an aliquot removed from the same material that was dosed in vivo) were also evaluated for internal structure by SAXS and cryo-TEM, and for apparent pKa using the TNS assay.

Luminescence signals were detected from the abdominal region of mice as early as 1-hour post injection for all samples and peaked at 3-6 h before decreasing to minimal levels by 48 h, consistent with previous reports (**Figure 4a**).^36,38^ Luminescence signal vs. time area under the curve (AUC) values were calculated over five timepoints then normalized to the hand mixing group within each study for combined comparison across both datasets (**Figure 4b**). All 1:1 FRR mixers (MIVM 1:1, Helix, CIJ) generated the weakest luminescence signals, again highlighting the importance of a second ‘quench’ mixing step immediately downstream of primary mixing to trap LNPs in a kinetic state during self-assembly. Turbulent mixing at a 3:1 FRR produced the strongest biological activity *in vivo*. LNPs made using the FDmiX yielded the highest radiance intensity, 4.6-fold higher than hand mixing. FDmiX claims the fastest mixing time (<1 ms) of all the devices tested, and results suggest that this geometry produces LNPs that outperform the others, *ceteris paribus*. Of the samples prepared at a 3:1 FRR, the Ignite and hand mixing produced LNPs that generated significantly lower *in vivo* luminescence than the other mixing strategies.

The *in vivo* results did not follow the same trend as the *in vitro* results, consistent with prior reports on poor *in vitro-in vivo* correlation.^39–41^ This highlights the limitations of *in vitro* screening of LNP development and emphasizes the importance of *in vivo* validation. For *in vitro* results, we expected smaller LNPs with lower PDIs and high EE values to show higher cellular uptake due to more efficient endocytosis, leading to stronger luminescence. However, statistical analysis revealed weak and inconsistent correlation between LNP physicochemical properties and mRNA translation activity, with neither Pearson nor Spearman tests identifying any single parameter as a reliable predictor (**Figure 3c-d**). *In vivo* efficacy (luminescence-time AUC) was correlated with particle size (r = –0.85, ρ = –0.93) and with encapsulation efficiency (r = 0.80, ρ = 0.77) (**Figure 4d-e**). A strong negative correlation was also observed between particle size and encapsulation efficiency (r = –0.98, ρ = –0.85), stronger than the pre-dialyzed samples, indicating that smaller particles consistently achieved higher cargo encapsulation and that became more pronounced after internal structural rearrangement through neutralization and ethanol removal.

Bioluminescence signal also decreased with increasing particle size (r = –0.85, ρ = –0.93). Apparent pKa exhibited a modest positive correlation with particle size (r = +0.48, ρ = +0.54), and modest negative correlation with *in vivo* signal (r = –0.47, ρ = –0.48). Mechanistically, this reflects the balance required for optimal pKa, because when pKa values exceed the 6.2–6.5 range, excessive surface protonation at physiological pH is implicated in impaired delivery.^42–44^ Zeta potential showed little correlation with either EE% or radiance (all r, ρ < ±0.3).

### Structural and morphological analysis

Small-angle X-ray scattering (SAXS) was used to evaluate internal structural features of the LNPs, as such data can be used in the development of structure-function relationships (**Figure 5a**).^45^ As with other physicochemical analyses in this work, notable differences were observed in SAXS profiles for LNPs prepared by different mixers – in this case, the location of the main peak (corresponding to internal structural spacings from 4.72 nm to 5.51 nm) and presence or absence of peaks at lower q, in the Porod region. We quantified the differences in scattering curves using the Volatility of Ratio (VoR) metric shown in **Figure 5b**. The VoR method compares SAXS intensity profiles at q-range of 0.2 – 1.8 nm^-1^ and reports the degree of divergence between them. A higher VoR value indicates more difference between two scattering patterns, which suggests structural variability between the two populations analyzed. Lower VoR values signify greater similarity.

**Figure 5.**
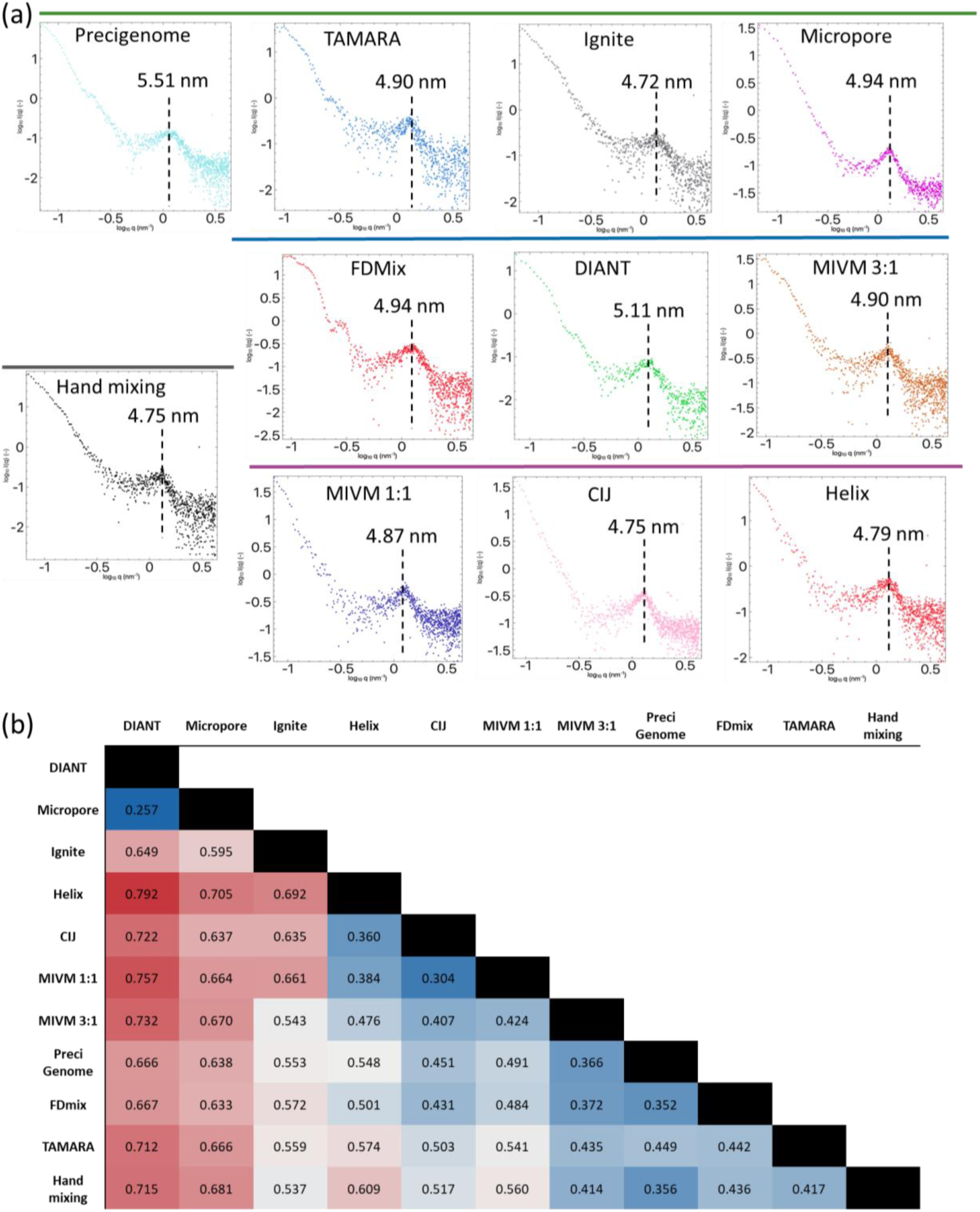
(a) SAXS scattering profiles of FLuc mRNA LNPs prepared with different mixers. Both axes are shown on a logarithmic scale, with the x-axis representing the scattering vector *q* (nm⁻¹) and the y-axis representing intensity I(q). Dashed lines mark the repeating unit distance within the nanoparticle internal structure. Colored lines above each chart denote the operating flow regime of the mixers: grey (undefined at 3:1 FRR), green (laminar at 3:1 FRR), blue (turbulent at 3:1 FRR), and purple (turbulent at 2:2 FRR).; (b) Volatility ratio (VoR) across mixers. Red boxes denote higher volatility values, indicating greater divergence between two mixers. Blue boxes denote lower values, reflecting internal structure similarity.

LNPs made by the DIANT mixer showed the greatest similarity to Micropore (volatility ratio of 0.257; same statistical group *in vivo*) and the highest divergence from Helix (0.792; different statistical group *in vivo*). The Ignite mixer produced LNPs with different internal structures from any other mixers, showing volatility values above 0.5 in most comparisons. The analysis shows comparability between the three mixers operated at a 1:1 FRR: the Helix shows strong similarity to CIJ (0.360) and MIVM 2:2 (0.407), and the CIJ was likewise similar closer to the MIVM 1:1 (0.384).

Cryo-TEM images showed that different mixing techniques can produce LNPs with varying morphologies as shown in **Figure 6**, consistent with previous work.^46,47^ The most pronounced differences were those between particles produced at a 3:1 and 1:1 FRR. Despite operating under different flow regimes, all 3:1 mixers yielded unilamellar, spherical, electron-dense particles. Precigenome and TAMARA mixers produced particles with more faceted morphologies than other systems. Particles produced using the Ignite mixer, on the other hand, exhibited numerous bleb-like protrusions. Importantly, all blebs observed in this study can be considered “empty” blebs, that do not display the characteristic mottled structure, indicative of mRNA partitioning into the bleb.^48^ Micropore, FDmiX, DIANT and MIVM 3:1 all generated uniform, spherical particles with electron dense cores. Interestingly, Helix (1:1 FRR) produced small, spherical particles with fewer empty vesicles and blebs, which is more similar to 3:1 mixers. However, cryo-EM micrographs represent a snapshot of the overall particle population, which does not reflect the particle size distributions observed in DLS.

**Figure 6.**
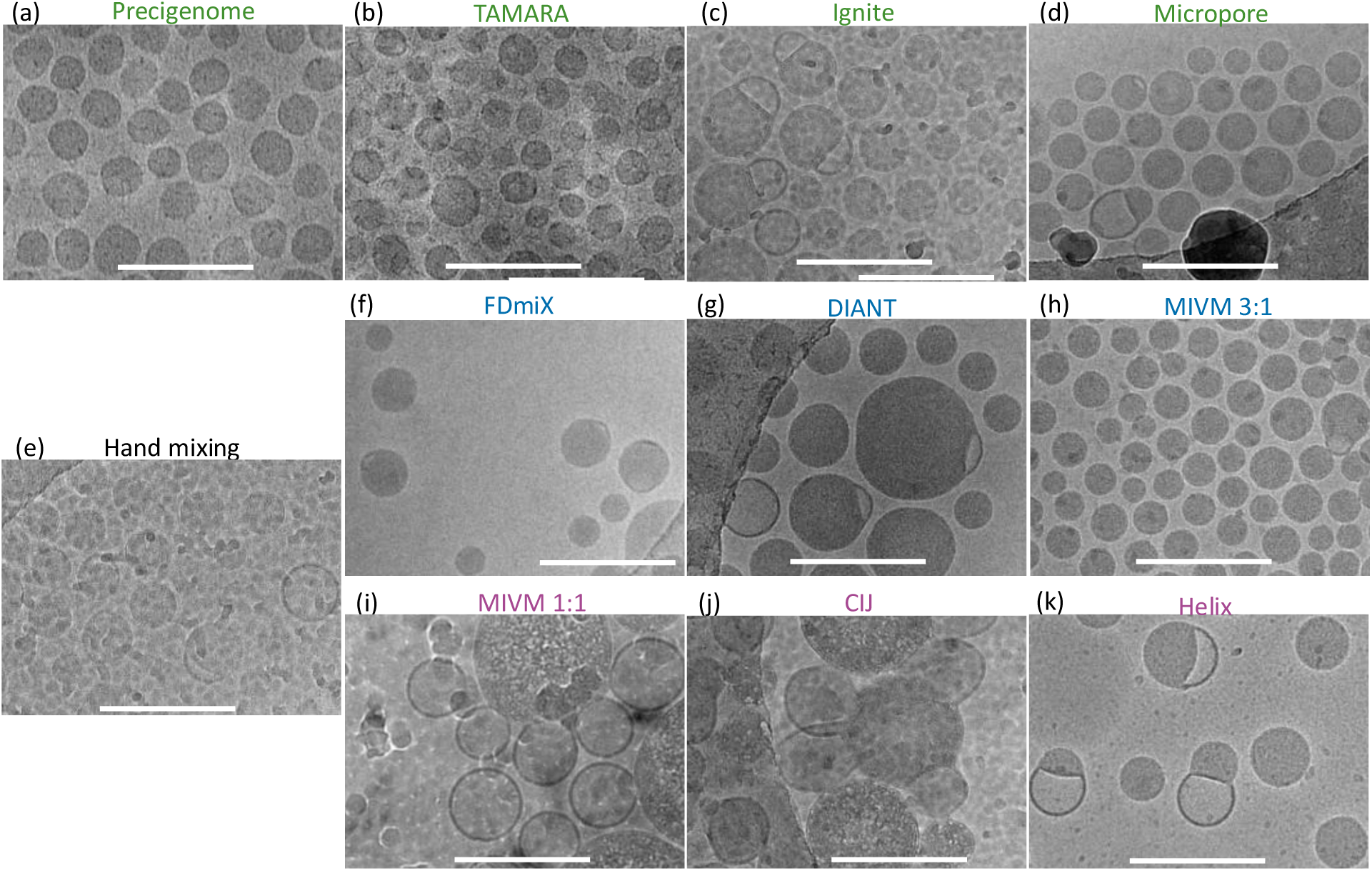
Cryo-electron microscopy (Cryo-EM) images of LNPs produced by different mixers. Green text denotes laminar mixers (3:1 FRR), blue text denotes turbulent mixers (3:1 FRR), and purple text denotes turbulent mixers (1:1 FRR). White scale bar represents 200 nm.

MIVM 1:1 and CIJ yielded large and heterogeneous vesicular structures, many of which appeared partially empty or membrane-bound with lower core contrast. The larger particles appear to be encapsulating the majority of the mRNA in this heterogenous population. These morphological features suggest fusion events, likely caused from multiple contributing factors. First, exposure to higher ethanol content prior to quenching enhances the outer lipid bilayer fluidity, making fusion energetically favorable.^49^ Post-processing steps, such as charge neutralization and concentration effects, can further weaken electrostatic stabilization, allowing vesicles to adhere and merge.^50^

### Conclusions

The goal of this study was not to identify one ‘best’ mixer but to demonstrate that the mixing technique used to prepare LNPs leaves significant and indelible fingerprints on the critical quality attributes of the resulting nanoparticles that persist through processing steps such as neutralization and concentration. If the population-level characterization techniques used here show such a significant degree of difference, the variability between particles within a population – e.g. spatial distribution of lipids in individual particles, distribution of lipid and RNA components across LNPs in a population – is also likely to differ significantly between materials made with differing mixing technologies, and more robust characterization using single-particle techniques is needed to study this in the future.

This point is particularly relevant as the use cases for LNPs become more complex and materials are being designed for e.g. protein replacement therapies, gene editing (involving co-loading of RNAs of multiple lengths into one particle), and extrahepatic delivery. Many of these next-generation delivery cases rely on highly-specific quality attributes, such as a surface chemistry that selectively binds one type of protein to affect biodistribution. Subtle differences in the arrangement and presentation of lipid components within and on the surface of LNPs in a population, which may be caused by differential mixing performance, are critical in such applications. There is a significant risk that promising biological performance seen from materials made using non-scalable mixing is due to critical quality attributes (CQAs) that were influenced by the specific mixing technique used, meaning those same CQAs will be different when the material is re-made using scalable mixing, and findings will not be generalizable or reproducible.

Mixing cannot remain an afterthought for this field. The mixing technique is an indispensable aspect of LNP formulation that must be considered explicitly. We therefore recommend that the field no longer consider as acceptable studies that provide no or only limited details about the specific mixing conditions used to prepare LNPs. Studies that utilize pipette-based manual mixing should be discouraged. Lessons learned from materials made using non-scalable microfluidic mixers should be further scrutinized and not considered generalizable unless demonstrated using LNPs made via more scalable techniques. To facilitate this transition in methods, manufacturers of large-scale mixing technologies should develop and validate equivalency of scaled-down versions of their mixers that can be operated with material consumption requirements comparable to the microfluidic mixers. For example, the ability of the MIVM to produce the same material whether operated with small-volume syringes or continuous pumps is an advantage – but the smallest feasible batch from the device used here requires 165 ug of RNA. Smaller-scale versions of scalable mixing geometries, or other methods to produce material that is representative of large-scale operation with minimal material consumption, should be developed, to facilitate research into the full relationship between mixing and LNP self-assembly.

## Methods

### Materials

ALC-0315 (6-((2-hexyldecanoyl)oxy)-N-(6-((2-hexyldecanoyl)oxy)hexyl)-N-(4-hydroxybutyl)hexan-1-aminium), DSPC (1,2-distearoyl-sn-glycero-3-phosphocholine), cholesterol, and DMG-PEG2000 (1,2-dimyristoyl-rac-glycero-3-methoxypolyethylene glycol-2000) were purchased from Avanti Polar Lipids (Alabama, USA). Purified Torulla Ambion yeast RNA (yRNA) was purchased from ThermoFisher Scientific (Massachusetts, USA). CleanCap® Firefly Luciferase (FLuc) mRNA containing unmodified nucleotides was purchased from TriLink Biotechnologies (California, USA). Free Acid OmniPur® HEPES was purchased from Fisher Scientific (New Hampshire, USA). Glacial acetic acid was purchased by Fisher Chemical. Anhydrous sodium acetate, ethanol 200 proof (100%), along with RNase-free water (Diethyl pyrocarbonate (DEPC) treated), were purchased from Fisher BioReagents (Pennsylvania, USA). For the in vitro study, D-Luciferin potassium salt was purchased from GoldBio (St. Louis, MO, USA). Fifty percent dextrose (Injection, USP) was purchased from Hospira (Lake Forest, IL, USA).

### Formulation and process parameters for lipid nanoparticle production

The organic feed solution was prepared by mixing ethanolic stock solutions of each individual lipid component to achieve a mixture of ALC-0315, DSPC, cholesterol, and DMG-PEG2000 at a molar ratio of 50:38.5:10:1.5 and a total lipid concentration of 6 mg/mL in ethanol. The acidic aqueous feed solution contained nucleic acid cargo (yeast RNA or FLuc mRNA) dissolved in pH 5.0 acetate buffer; 20 mM buffer and 0.3 mg/mL RNA for mixers operated at a flowrate ratio (FRR) of 1:1, and 13.32 mM buffer and 0.1 mg/mL RNA for mixers operated at 3:1 FRR, such that the ionic strength at the point of mixing was identical across all runs. Organic and aqueous feed solutions were combined using each mixing technology at the operational FRRs and total flow rates (TFRs) provided in **Table S1**. Collected efluent mixtures were quenched (diluted) to 10% v/v ethanol to reduce Ostwald ripening using 10 mM acetate buffer (pH 5.0) dispensed into the collection vial by pipette 5 seconds after a run was stopped, to allow LNPs to completely form at the conditions in the mixing chamber before subsequent dilution. The final composition of all LNP suspensions produced in this way was identical across all mixers: 0.6 mg/mL lipids, nucleic acid concentration of 0.03 mg/mL, an N/P ratio of 5:1 (cationic ionizable lipid: nucleic acid) and a lipid-to-nucleic acid mass ratio of 20:1, in 90/10 v/v 10mM acetate pH 5 / ethanol.

To remove residual ethanol and raise pH to neutral, FLuc mRNA@LNP formulations were dialyzed against 10 mM HEPES buffer (pH 7.4) using Spectra Por S/P 1 Dialysis Membrane (6-8 kDa MWCO) for 4 hours. The dialyzed LNP suspensions were concentrated, using centrifugal filtration, to a total lipid concentration of 10–18 mg/mL using Amicon Ultra centrifugal filters (100 kDa MWCO, Millipore Sigma) on a centrifuge operated at 500 rcf. Prior to concentration, the filters were primed via centrifugation at 500 rcf in three 2 min cycles. The first priming used a mixture of water and ethanol (70:30), followed by two separate rinsing cycles with RNase-free water. The concentration factor for each suspension was calculated using the Ribogreen assay to calculate the total concentration of RNA in a sample.

### Mixing Platforms

Hand mixing: Eppendorf Research™ plus, Mechanical Single-Channel Pipettes (Hamburg, Germany. Cat no. 3123000063) were utilized to dispense the organic lipid solution into the aqueous solution at a 1:3 volumetric ratio. After dispensing the organic feed, fifty rapid aspiration–dispense cycles were performed to mix the fluids.

NanoAssemblr™ Ignite Mixer: The NanoAssemblr™ Ignite system (Ontario, Canada) employs a toroidal microstructure in the NxGen™ micromixer single-use cartridge constructed from polymethyl methacrylate (PMMA). The channel depth for all pathways is 290 μm, with a width of 80 μm in the main mixing regions. LNPs were prepared at a TFR of 12 mL/minute, using a of 3:1 FRR dispensed from BD Luer-Lok™ syringes using the integrated console pump. An initial volume of 0.3 mL was diverted to waste prior to sample collection.

PreciGenome Nanogenerator: PreciGenome Nanogenerator CHP-MIX-4 (California, USA) employs a Tesla microstructure fabricated by polymer injection molding. Each chip contains a four-tesla mixing line that can generate LNP volumes ranging from 250 µL to 3 mL. The chips’ inlets and flow paths were primed at the experimental flow rates for 45 seconds before sample collection. The mixer was then operated at a TFR of 4 mL/min with an aqueous-to-organic FRR of 3:1, using 3 mL BD Luer-Lok™ syringes driven by syringe pumps (Harvard Apparatus, Massachusetts, USA).

TAMARA Mixer: The TAMARA InsideTX chip (InsideTX, Paris, France) uses a staggered herringbone structure for LNP production. The TAMARA system consists of a control module and a mixing module. The mixing module delivered the feed streams to the chip using compressed nitrogen at a pressure of 8 bar. Feed solutions were pipetted into the liquid reservoirs of the mixing module and mixing was carried out at a TFR of 5 mL/min and a 3:1 FRR.

FDmiX by Fluid Dynamix: The FDmiX M mixer (Berlin, Germany) is made of stainless steel and incorporates an OsciJet nozzle that generates high-frequency oscillatory mixing. The mixer was operated at a TFR of 80 mL/min with two inlets operating at a 3:1 FRR. Before sample collection, inlets and flow paths were primed at the experimental flow rates for 30 seconds. Feed solutions were delivered using 30 mL BD Luer-Lok™ syringes driven by syringe pumps.

LARU Discovery by DIANT: The LARU Discovery system (DIANT, Massachusetts, USA) was operated at a TFR of 80 mL/min using a 3:1 aqueous-to-organic FRR. An initial volume of 11 mL was discarded as waste prior to LNP sample collection. Feed solutions were delivered to the shearing co-axial jets mixer using the system’s integrated syringe pump.

AXF Mini by Micropore Technologies: The AXF mini (Micropore Technologies, Teesside, UK) was primed at the experimental flow rate (80 mL/min) for 30 seconds prior to LNP production. The LNPs were prepared using a TFR of 80 mL/min and a 3:1 FRR.

Multi-Inlet Vortex Mixer (MIVM): The MIVM (Holland Applied Technologies, US) is a four-inlet mixer that leverages vortex-induced turbulence to combine feed streams at a total flow rate of 160 mL/min. This mixer was used twice, once at a 1:1 FRR and once at a 3:1 FRR, to interrogate the difference of solvent quality at the point of mixing while other parameters, including mixing geometry, were held constant. For operation at a 1:1 FRR, two inlets contained the organic phase, and two inlets contained 0.3 mg/mL nucleic acid in 20 mM acetate buffer at pH 5. For a 3:1 FRR, a single inlet contained the organic phase, one inlet contained an aqueous stream of 0.3 mg/mL nucleic acid in 13.32 mM acetate buffer pH 5, and the remaining two inlets contained streams of 13.32 mM acetate buffer at pH 5 without dissolved RNA. Based on previous experience, the organic and nucleic acid-containing aqueous streams were positioned opposite each other on the device^51^. Each inlet was loaded with 0.55 mL of feed solution in 1 mL NORM-JECT (Henke-Sass Wolf, Germany) syringes and manually depressed using a stainless steel base plate mixer stand (Holland Applied Technologies) to ensure equal velocity and pressure, which has been previously demonstrated to drive mixing equivalent to pump-driven flow at low total material consumption.^52–54^

Confined Impinging Jets Mixer (CIJ): The CIJ (Holland Applied Technologies, US) utilized in this work is made of Delrin plastic. 0.55mL of ethanol and buffer feed solutions were loaded into 1 mL NORM-JECT all-plastic luer-slip syringes that were manually depressed using equal pressure to achieve a total flow rate of 120 mL/min, a technique which has been previously demonstrated to drive mixing equivalent to pump-driven flow at much lower total material consumption.^52–54^

Nova Impinging Jets Mixer by Helix Biotech: The Nova Impinging Jets Mixer (Helix Biotech, Tennessee, US) is machined from stainless steel and also utilizes impinging jets to achieve homogenization of the inlet streams at a 1:1 FRR. Each inlet was loaded at equal volume into 3 mL syringes and dispensed at equal pressure using the built-in syringe pumps delivering flow at a TFR of 8 mL/min. A volume of 1.5 mL was diverted to waste automatically enabled by the in-line waste-collection line switcher prior to LNP sample collection.

### Physicochemical, Structural and Biological Characterization of LNPs

LNPs were characterized pre-dialysis, post-dialysis, and post-concentration for particle size, polydispersity, and zeta potential using dynamic light scattering (DLS) described below. Encapsulation efficiency (EE) was determined via the RiboGreen assay as described below. DLS and EE measurements on concentrated LNPs samples were performed after back-dilution by the corresponding concentration factor to enable direct comparison with pre-concentrated samples.

#### Physicochemical properties

Size, polydispersity index (PDI), and zeta potential (ζ) of LNP samples were measured using a Malvern Zetasizer Pro (Malvern Panalytical, UK). For size and PDI measurements, LNP samples were diluted 1:10 in their respective buffer systems (10 mM acetate buffer pH 5 or 10 mM HEPES buffer pH 7.4) and transferred into 1.5 mL semi-micro polystyrene cuvettes (Fisherbrand, US). Following production, samples were left briefly at room temperature to degas trapped air that may be introduced by high-velocity mixing, and DLS measurements were then performed within minutes. Ζeta potential measurements were performed by diluting the LNP samples 1:10 in 20 mM NaCl and loading the diluted sample into a folded capillary zeta cell (Malvern Panalytical, UK). Each measurement consisted of three runs with 15 readings per run at 173° scattering angle.

#### Encapsulation efficiency

The mRNA encapsulation efficiency and absolute mRNA content were determined using the Quant-iT™ RiboGreen assay (Cat# R11490, Life Technologies). Working solutions of 0.5% Triton X-100 and 0.5% RiboGreen dye were prepared in Tris EDTA (TE) buffer. Samples treated with Triton X-100 represented total nucleic acid content (encapsulated + unencapsulated), and untreated samples represented unencapsulated RNA only. Calibration curves were generated by measuring fluorescence of standard RNA solutions (0, 0.1, 0.2, 0.4, 0.6, 0.8, and 1 µg/mL in TE buffer, pH 7) with and without Triton X-100, each containing 0.25% RiboGreen dye.

For sample analysis, LNP suspensions were diluted to 8 µg/mL RNA content using TE buffer. A volume of 26.25 µL of the LNP suspension was added to 148.75 of TE buffer with and without 2% Triton X-100 in parallel. To this mixture, a volume of 175 µL of 0.5% RiboGreen dye was added, and the mixture was left to incubate protected from light for 5 min. A 100 µL of each sample and calibrant mixtures were transferred to an opaque 96-well plate (black) for analysis. Calibration standards were prepared in duplicate whereas experimental samples were prepared in triplicate. Fluorescence intensities were recorded on a Molecular Device SpectraMax i3x Paradigm Multi-Mode Detection Platform (California, US), setting the λ_excitation_ = 485 nm and the λ_emission_ = 528 nm. RNA concentrations were determined by comparing sample fluorescence to the calibration curve gradient. Encapsulation efficiency was calculated according to the following formula:

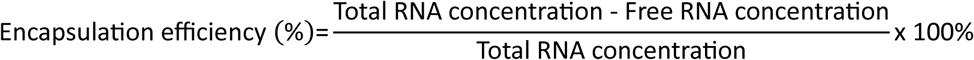

### Cell culture and transfection

HEK293 cells (ATCC, Manassas, VA, USA) were cultured in DMEM medium supplemented with 10% fetal bovine serum (FBS), 100 U/mL penicillin, and 100 μg/mL streptomycin at 37°C in a humidified atmosphere containing 5% CO_2_. For LNP transfection, HEK293 cells were seeded at 50,000 cells per well in 24-well plates and allowed to adhere for 24 hours. Cells were treated with FLuc mRNA-loaded LNPs and incubated for an additional 24 hours. After treatment, the cell culture medium was removed, and cells were rinsed with 1x PBS. Cells were then lysed in 150 µL of lysis buffer (Promega, Madison, WI, USA), and the lysate was transferred to a 1.5 mL microcentrifuge tube. Samples were centrifuged at 17,000 x g for 4 minutes to remove cellular debris. The protein content in the supernatant was determined by the Pierce BCA Protein Assay Kit (Thermo Fisher Scientific, Rockford, IL, USA) and used to normalize luminescence intensity. For measurement of luminescence intensity, which represents luciferase expression, 20 µL of cell lysate was mixed with 100 µL of Luciferase Assay Reagent (Promega, Madison, WI, USA) and analyzed with a BioTek Synergy Neo2 microplate reader (Agilent, Santa Clara, CA, USA). A dose response of mRNA transfection was examined to determine the mRNA concentration to treat HEK293 cells for the in vitro protein expression test (**Figure S2**). Unless specified otherwise, the cells were treated with FLuc mRNA-loaded LNPs at a total mRNA concentration (both encapsulated and unencapsulated mRNA in each preparation) of 0.65 µg/mL in the medium, the half-maximal effective concentration (EC50).

### Bioluminescence imaging of FLuc mRNA LNP Transfection

All animal procedures were approved by the Institutional Animal Care and Use Committee in conformity with the NIH guidelines for the care and use of laboratory animals. Five-six weeks old female BALB/c mice were purchased from Envigo (Indianapolis, IN, USA) and acclimatized for one week before the procedure.

BALB/c mice were individually weighed. FLuc mRNA-loaded lipid nanoparticles (LNPs), as diluted in 5% dextrose (D5W), were administered intravenously at a nominal total mRNA dose of 0.2 mg/kg. At designated time points (1, 3, 6, 24, and 48 hours post-injection), mice were intraperitoneally injected with D-luciferin (50 mg/kg). Five minutes after luciferin administration, bioluminescence imaging was performed using the Spectral Ami Optical Imaging System Spectral Instruments Imaging (Tucson, AZ, USA). Mice were anesthetized with 2.5% isoflurane and maintained under anesthesia in the supine position during imaging. Bioluminescent signals were quantified as radiance (photons/s/cm²/sr) from the abdominal region using Aura In Vivo Imaging Software, Spectral Instruments Imaging (Tucson, AZ, USA).

### Benchtop small angle X-ray scattering

Small-angle X-ray scattering (SAXS) experiments were carried out on a SAXSpoint 2.0 instrument (Anton Paar) with a copper Kα source (λ = 1.54 Å). Measurements were performed using an incident beam, fixed at 8.05 keV and collimated under a vacuum using an advanced scatterless beam collimation (Anton Paar). Sample-to-detector distance (SDD) was set to 575 mm, which corresponded to a q-range of 0.14-4.2 nm^-1^. An EIGER R series Hybrid Photon Counting detector was used for data recording. LNP samples or background buffer were pipetted into 1 mm quartz capillaries (Anton Paar). Data collection was carried out by averaging five frames of 360 s exposure time. The 2D scattering data were reduced using SAXSanalysis, Version 2.50 (Anton Paar). Background (buffer) subtraction was performed using the PRIMUS package from ATSAS 3.2.1.^55^

### Cryo-EM

Quantifoil® R 1.2/1.3 300 mesh Cu grids (Quantifoil, Germany) were glow-discharged for 60 seconds at 25mA on a PELCO easiGlow system (Ted Pella). A 3 µL aliquot of concentrated lipid nanoparticles was applied to the grids utilizing Vitrobot Mark IV (Thermo Fisher Scientific) and subsequently plunged frozen in liquid ethane. Grids were stored in liquid nitrogen until imaging. Cryo-transmission electron microscopy was performed on a Talos F200C with a Ceta 4k x 4k CMOS camera (Thermo Fisher Scientific).

### TNS Assay

The apparent pKa of LNPs was measured using the LipidLaunch™ LNP Apparent pKa Assay Kit TNS (6-(p-toluidino)-2-naphthalenesulfonicacid sodium salt) (Cayman Chemical, Ann Arbor, MI, USA). Prior to starting the assay, concentrated FLuc mRNA LNPs were diluted to a final lipid concentration of 6 mg/mL using 10 mM HEPES buffer pH 7.4.

The buffer was prepared by diluting the twelve 10x stock reagents supplied with the kit to a 1x concentration using RNase-free water. The resulting 1x buffers spanned a pH range of 4.0 to 9.0. The pH of each 1x buffer was measured to two decimal places using a Thermo Scientific Orion STARA1110 Star A111 Benchtop pH Meter (Massachusetts, USA) and Thermo Orion™ 9106BNWP pH probe (Massachusetts, USA). Before combining the buffer and the LNPs, the supplied TNS probe solution was diluted to 83µM using RNase-free water.

Each well in a black, flat-bottom Corning 96-well microplate (US) was loaded with 90 µL of 1x buffer, 5 µL of LNP sample, and 5 µL of diluted TNS probe solution to yield 12 wells per replicate (1 sample x 12 pH levels). Each condition was prepared in triplicate (36 wells per formulation). The prepared plate was covered and incubated on a plate shaker at room temperature for 20 minutes. The fluorescence intensity was recorded using Molecular Devices SpectraMax i3x Paradigm Multi-Mode Detection Platform (California, US) setting the λ_excitation_ = 320 nm and the λ_emission_ = 450 nm. Raw fluorescence readings were averaged across replicates, and the mean intensity of the pH 9.0 buffer was subtracted from all values to yield the average relative fluorescence (ARF). The apparent pKa was calculated by fitting a four-parameter logistic (4PL) regression model as shown in equation below:

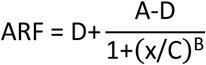

where A, B, C, and D are fitting constants, and C corresponds to the pH value of half-maximal intensity (apparent pKa). Curve fitting was performed in MATLAB (R2024, Mathworks, Natick, MA, USA) using the 4PL regression code^56^. Fittings shown in **Figure S3**.

### Statistical analysis

Statistical analyses on mixers producing LNPs encapsulating yeast RNA were performed using ordinary one-way ANOVA with a 95% confidence level. Post-hoc comparisons were conducted using Tukey’s HSD test, and pairwise differences were summarized through compact letter displays. Pearson correlation coefficients (r) and Spearman rank correlations (ρ) were calculated to quantify the strength of linear relationships and monotonic trends, respectively. Ordinary one-way ANOVA analysis was carried out using GraphPad Prism and correlation coefficient calculations were done using Microsoft Excel.

## Acknowledgements

This research was supported by a grant from Eli Lilly and Company (USA), part of the Eli Lilly and Purdue University Research Alliance Center (LPRC). The authors would like to posthumously thank Dr. Young-Ho Song at Eli Lilly for her insightful comments and advice over the course of the study’s development and execution.

We thank Dr. Bernhard Bobusch from Fluid Dynamix, Dr. Camden Cutright from Micropore Technologies, and Dr. Robin Oliveres from InsideTx for loaning mixers to be used in the study, and Dr. Tony Costa from DIANT for insightful discussions about the operation of the LARU mixer.

Cryo-EM data were obtained in the Purdue University Bindley Bioscience Center. C. Sams (Agricultural and Biological Engineering, Purdue University) assisted with LNP preparation and data visualization. C. French (Biomedical Engineering, Texas A&M University) assisted with *in vivo* tests. We thank the Purdue Imaging Facility for their assistance with IVIS imaging and data collection.

## Author contributions

TB – methodology, investigation, formal analysis, validation, data curation, writing – original draft preparation, visualization

AA – investigation, writing – reviewing and editing

LY – investigation, writing – original draft preparation HM – investigation, writing – original draft preparation MB – investigation, writing – reviewing and editing

GH – investigation, writing – reviewing and editing SRD – investigation, writing – reviewing and editing FM – investigation, writing – reviewing

FV – investigation, writing – reviewing and editing

SLH – investigation, writing – original draft preparation

MF, LAM, PV, AA, YY, KDR – conceptualization, methodology, project administration, funding acquisition, supervision, writing – reviewing and editing.

All authors have read and agreed to the published version of the manuscript.

