## Supporting information for "Identifying differential effects from eleven mixing techniques on mRNA lipid nanoparticle physicochemistry and biological performance"

Table of contents and figures:


- Discussion: descriptions of mixers
- Figure S1: workflow
- Table S1. Operating Conditions of Mixers
- Figure S2: LNP heatmaps with standard deviations
- Figure S3: dose-response curve for in vitro dosing
- Discussion: in vivo dosing
- Discussion: TNS assay
- Figure S4: TNS assay (apparent pKa) fitting for each mixer

**Discussion: descriptions of mixers**

Hand mixing is designated as the base case in this study. It relies on the mechanism of a spring inside the pipette and the speed of the aspirate-dispense cycle to drive mixing and nanoprecipitation, forming lipid nanoparticles with little to no holdup volume in the system. Hand mixing is user dependent. While its internal reproducibility is satisfactory, its external reproducibility remains a challenge. [18,20] The same user prepared all hand-mixed samples in this study.

PreciGenome, TAMARA, and Ignite are passive microfluidic mixers that have microstructures of tesla valves, staggered herringbone, and toroidal rings, respectively, shown with green bars in the graphical abstract. Modern micromixers rely on channel geometry for mixing through a synergetic mechanism of diffusion and chaotic advection, aiming to increase the contact surface area between organics and aqueous feed solutions. These laminar mixers have small channels which facilitate low velocity of fluid; thus, the fluids exhibit low Reynolds numbers in the mixing chamber. PreciGenome (tesla) and Ignite (toroidal) are employed as representatives of mixers that take advantage of their geometry of turns to repeatedly disturb the laminar flow, creating micro-vortices, and achieve homogeneous mixing, while TAMARA utilizes a ribbed pattern to induce chaotic mixing[26].

PreciGenome employs a Tesla microstructure which features a sequence of asymmetrical channels. It generates varying flow resistance as the fluid advances, splitting into branches that rejoin at different velocities and directions[26]. With each circulation, these disturbances are amplified and leading to rapid homogenization despite low operating flow rate of 4 mL/min and thus has a relatively long residence time. This low flow rate may allow more delicate control that dictates the nanoparticle characteristics as shown in main body **Figure 1**, exhibiting the tightest size distribution (polydispersity index), smallest zeta potential, and highest encapsulation efficiency.

TAMARA is a platform that allows the option of either herringbone or baffle microstructure as both were embedded in a single chip, and in this study we used its staggered herringbone microstructure. It requires 8 – 10 barg supplied by compressed nitrogen to generate a pressure gradient allowing organics and aqueous streams to flow through the mixing geometry. The intuitive interface of TAMARA facilitates a pressure test to check whether the platform can receive and contain high pressure to assure lipid nanoparticle production is at the optimal condition. Proper cleansing of chips using water and ethanol enabled up to 10 uses per chip, with the cleaning procedure programmed on its console effectively after lipid nanoparticle production, by air drying, water-ethanol mixing, followed by air drying.

The NanoAssemblr™ Ignite chip is the most common microfluidic mixers for lipid chemistry exploratory study and LNP production in academics[6–8,27–29]. The Ignite chip employs a toroidal microstructure in which the aqueous and organic streams are guided through concentric ring channels that fold and stretch the interface multiple times. This mechanism enhances diffusion to promote LNP self-assembly.[30]

The AXF Mini produced by MicroPore operates in the laminar flow regime but its physical dimensions are larger than what is typically considered microfluidic. The device is a compact crossflow membrane mixing system which places the organic stream as a dispersed phase that is pushed through the membrane pores where it breaks into fine droplets. These droplets immediately interact with the aqueous stream which serves as the continuous phase flowing tangentially along the membrane surface. This tangential crossflow keeps the membrane clear and prevents clogging by sweeping away newly formed droplets while maintaining a consistent shear at the pore openings. An optional integrated computer interface is available to procure along with the AXF Mini mixer, but the mixer alone can be operated with third-party syringe pumps.

Turbulent mixers operate under flow regimes where the Reynolds number is sufficiently high to generate chaotic eddies and vortices that dominate over molecular diffusion. FDmiX, DIANT, MIVM, CIJ, and Helix are categorized as turbulent mixers because these mixers accelerate fluid streams at high velocities through constricted geometries, such that inertial and convective forces outweigh viscous damping. Then, vigorous turbulence that promotes rapid homogenization of organic and aqueous phases occurs. These mixers are depicted with purple bars in main body **Figure 1**.

FDmiX incorporates an OsciJet nozzle that generates high-frequency oscillatory mixing by repeatedly deflecting the fluid jet of primary stream (aqueous) from side to side within the main chamber. As the primary stream oscillates, a portion of the jet also is diverted through a side channel and reintroduced to drive the next deflection, creating continuous jet switching. This oscillatory motion enhances turbulence and promotes rapid interaction with a secondary stream (organics) which is introduced perpendicularly at the nozzle inlet. Within the mixing chamber, the oscillating jet impinges on the perpendicular stream, resulting in intense mixing before the combined flow exits the outlet.[31]

DIANT LARU is a coaxial mixer that incorporates two concentric tubes, with the organic solution injected in the same direction as the surrounding aqueous solution. The organic stream flows through the inner tube, and the aqueous stream flows through the outer tube, such that the two meet at the outlet of the inner tube. When Reynolds numbers are high, the sharp velocity gradient at the fluid interface creates strong shear layers that quickly evolve into turbulence. In this regime, swirling vortices and eddies continuously distort the liquid interface, enhancing convective mixing and driving faster blending between two streams. The outlet tube includes a flow splitter, which supplements the built-in line switcher connected to the mixer console, automating switching between waste and sample collection. Syringe size, inlet flow rate, and flow rate ratio, pre-waste volume, and dilution factor information are modulated via DIANT® software through a USB connection. An optional second mixing point is included in the device, where a peristaltic pump can be used to dilute liquid from the primary mixer. This feature was not used in this study, in order to isolate the effect of primary mixing.

The Multi-Inlet Vortex Mixer (MIVM) employs the Flash NanoPrecipitation (FNP) technique to generate nanoparticles with high reproducibility. Its design incorporates four independent inlets, enabling flexible combinations of feed orientations, phases, and components, while allowing each stream's flow rate to be controlled individually to enhance micromixing.[32] This versatility makes the system adaptable to diverse formulations and user-friendly, as it can be operated either manually or with syringe pumps at efficient total flow rates up to 160 mL/min. In this study, equal volumes of organic and aqueous streams were introduced tangentially into the cylindrical chamber of the MIVM using syringes pushed with a horizontal plate on rails, as demonstrated previously. The resulting vortex repeatedly folded and stretched the liquid interfaces within milliseconds, leading to LNP formation.

The Confined Impinging Jets (CIJ) mixer employs two opposing inlets instead of the MIVM's four. In this configuration, high-velocity organic and aqueous streams collide head-on in a central chamber, producing intense turbulence that drives nanoprecipitation through the Flash NanoPrecipitation (FNP) technique. The CIJ retains the same flexibility as the MIVM, with operation possible by either manual syringe injection or pump-driven flow at total flow rates of 120 mL/min.

Similar to the CIJ, Helix (Nova™ IJM) also employs impinging jet technology, with interchangeable mixers of different sizes to match desired throughput, ranging from 8 mL to 122 mL. In this study, Helix was operated with its smallest mixer compartment as a scaled-down analogue of the CIJ, at a total flow rate of 8 mL/min. The system is composed of three main consoles: one for the syringe pump, one for the mixer, and one for holding the collection and waste vials. An automated valve switcher directs the output between waste and collection vials which enables convenient sample collection. Like with the DIANT LARU, an optional second mixing point can be purchased with the device, to dilute liquid from the primary mixer shortly after primary mixing. This feature was not used in this study.

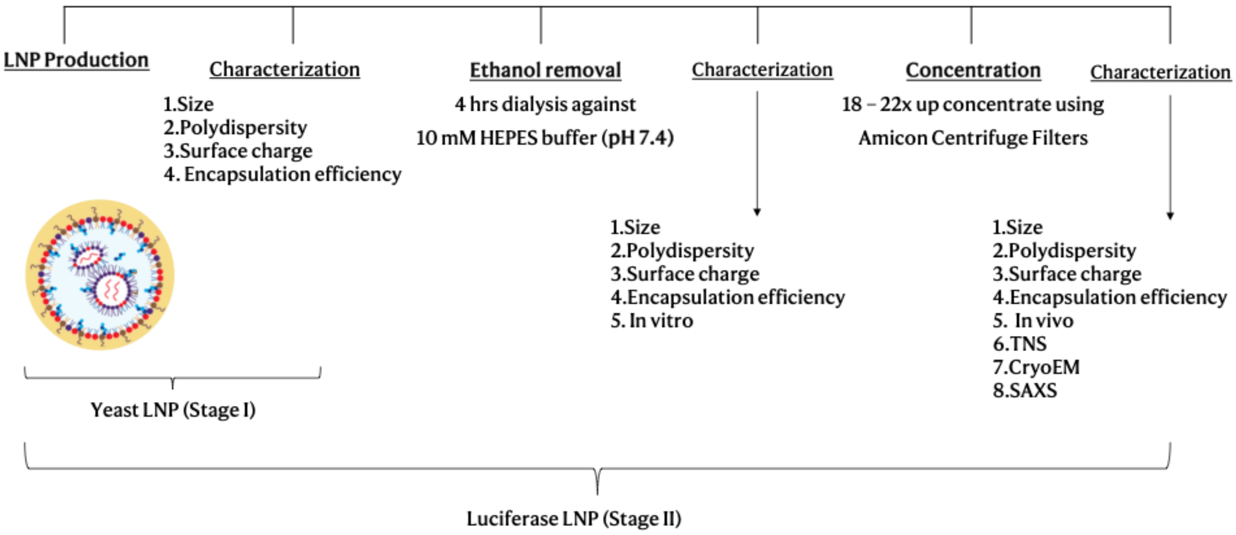


**Figure S 1.** Schematic representation of the experimental workflow used for the production and downstream processing and characterization of yeast and luciferase mRNA lipid nanoparticles.

**Table S1. Operating Conditions of Mixers**

**Table S1.** Operating Conditions of Mixers

| Mixer | Mixing technology | Inlets operation | Aqueous-to-Organic FRR | TFR (mL/min) |
|---|---|---|---|---|
| **Hand mixing** | Pipette | Manual | 3:1 | - |
| **PreciGenome Nanogenerator (PreciGenome)** | Tesla geometry (microfluidic) | Harvard Apparatus syringe pump | 3:1 | 4 |
| **TAMARA InsideTx (TAMARA)** | Herringbone geometry (microfluidic) | Compressed nitrogen | 3:1 | 5 |
| **Nano-Assemblr® Ignite™ Precision Nanosystems (Ignite)** | Toroidal geometry (microfluidic) | Built-in system syringe pump | 3:1 | 12 |
| **FDmiX M (Fluid Dynamix)** | Oscillatory | Harvard Apparatus syringe pump | 3:1 | 80 |
| **LARU Discovery (DIANT)** | Coaxial | Built-in system syringe pump | 3:1 | 80 |
| **AXF Mini (Micropore)** | Cross-flow | Harvard Apparatus syringe pump | 3:1 | 80 |
| **Multi-Inlet Vortex Mixer (MIVM)** | Vortex | Manual | 3:1 and 1:1 | ≈160 |
| **Nova™ IJM (Helix Biotech)** | Impinging jets | Built-in system syringe pump | 1:1 | 8 |
| **Confined Impinging Jets (CIJ)** | Impinging jets | Manual | 1:1 | ≈120 |

**Figure S2: LNP heatmaps with standard deviations**

| Mixers | Size (nm) | PDI | Zeta Potential (mV) | EE% |
|---|---|---|---|---|
| Ignite | 44.83± 2.34 | 0.26± 0.01 | 4.27± 0.46 | 86.56± 0.79 |
| DIANT | 47.16± 3.33 | 0.26± 0.01 | 3.31± 1.09 | 91.91± 0.60 |
| TAMARA | 61.48± 2.26 | 0.21± 0.01 | 6.01± 1.15 | 85.02± 1.61 |
| MIVM 3:1 | 63.42± 4.78 | 0.23± 0.02 | 5.88± 0.49 | 79.71± 2.72 |
| FDmiX | 63.58± 1.53 | 0.20± 0.02 | 3.52± 0.33 | 88.46± 0.93 |
| Precigenome | 74.16± 0.99 | 0.08± 0.01 | 1.12± 0.88 | 94.63± 1.88 |
| Micropore | 74.56± 1.14 | 0.19± 0.02 | 1.96± 0.81 | 76.37± 2.29 |
| CIJ | 76.39± 4.86 | 0.16± 0.03 | 5.64± 0.99 | 67.17± 2.89 |
| Hand mixing | 80.07± 5.76 | 0.16± 0.02 | 5.63± 0.84 | 90.58± 1.16 |
| Helix | 92.89± 1.65 | 0.15± 0.02 | 2.88± 1.46 | 74.99± 3.56 |
| MIVM 1:1 | 103.71 ± 2.01 | 0.21± 0.01 | 3.58± 1.13 | 73.85± 1.09 |

**Figure S 2**. Yeast LNP heatmap with standard deviation extending Figure 1e

| Mixer | Size (nm) | PDI | Zeta Potential (mV) | EE% |
|---|---|---|---|---|
| Ignite | 63.35 ± 0.98 | 0.23 ± 0.02 | 8.19 ± 2.21 | 95.95 ± 0.03 |
| FDmiX | 64.56 ± 0.42 | 0.09 ± 0.02 | 2.36 ± 1.74 | 85.20 ± 0.10 |
| TAMARA | 68.12 ± 0.54 | 0.17 ± 0.01 | 3.20 ± 1.12 | 91.13 ± 0.50 |
| DIANT | 69.07 ± 0.78 | 0.17 ± 0.03 | 3.22 ± 1.55 | 91.70 ± 0.03 |
| Precigenome | 72.80 ± 0.44 | 0.10 ± 0.04 | 1.20 ± 0.67 | 93.34 ± 0.08 |
| Micropore | 77.16 ± 0.96 | 0.09 ± 0.03 | 0.87 ± 0.78 | 95.16 ± 0.28 |
| MIVM 3:1 | 77.65 ± 1.25 | 0.33 ± 0.01 | 16.97 ± 0.48 | 93.06 ± 0.09 |
| Hand mixing | 96.34 ± 1.46 | 0.15 ± 0.02 | 3.35 ± 0.30 | 86.98 ± 0.89 |
| Helix | 99.33 ± 0.61 | 0.15 ± 0.04 | 6.20 ± 0.68 | 55.56 ± 1.09 |
| CIJ | 102.00 ± 1.27 | 0.13 ± 0.01 | 5.35 ± 0.44 | 68.51 ± 1.79 |
| MIVM 1:1 | 115.00 ± 1.08 | 0.15 ± 0.02 | 7.48 ± 0.68 | 59.81 ± 1.81 |

**Figure S 3**. Fresh FLuc LNP heatmap with standard deviation extending Figure 2d

| Mixer | Size (nm) | PDI | Zeta Potential (mV) | EE% | Luminescence (% relative to HM) |
|---|---|---|---|---|---|
| FDmiX | 67.48 ± 0.99 | 0.09 ± 0.05 | -4.80 ± 1.42 | 96.92 ± 0.05 | 225.25 ± 6.83 |
| DIANT | 74.70 ± 0.67 | 0.09 ± 0.01 | -4.62 ± 3.84 | 96.27 ± 0.18 | 169.72 ± 24.23 |
| TAMARA | 77.77 ± 0.30 | 0.19 ± 0.01 | -0.70 ± 0.17 | 90.13 ± 0.51 | 217.80 ± 18.13 |
| Micropore | 84.57 ± 1.41 | 0.06 ± 0.02 | -2.64 ± 3.45 | 93.95 ± 0.06 | 462.95 ± 31.26 |
| Ignite | 85.84 ± 8.05 | 0.22 ± 0.01 | -0.15 ± 0.68 | 85.69 ± 0.77 | 187.74 ± 12.47 |
| Precigenome | 90.29 ± 0.55 | 0.07 ± 0.01 | -12.55 ± 3.71 | 86.63 ± 0.97 | 162.99 ± 46.86 |
| MIVM 3:1 | 93.14 ± 1.25 | 0.34 ± 0.04 | 0.04 ± 0.34 | 91.54 ± 1.53 | 35.15 ± 4.56 |
| Hand mixing | 99.06 ± 1.06 | 0.18 ± 0.02 | -0.38 ± 2.00 | 95.39 ± 0.15 | 100.00 ± 3.03 |
| CIJ | 108.80 ± 1.84 | 0.09 ± 0.02 | -4.60 ± 1.39 | 71.86 ± 1.75 | 328.01 ± 18.37 |
| Helix | 109.30 ± 0.19 | 0.09 ± 0.00 | -5.70 ± 1.53 | 69.52 ± 0.42 | 97.02 ± 26.26 |
| MIVM 1:1 | 135.90 ± 1.08 | 0.17 ± 0.03 | -1.48 ± 1.65 | 57.11 ± 2.43 | 517.96 ± 6.87 |

**Figure S 4**. Dialyzed FLuc LNP heatmap with standard deviation extending Figure 3

| Mixer | Size (nm) | PDI | Zeta Potential (mV) | EE% | Radiance AUC (% relative to HM) | Apparent pKa |
|---|---|---|---|---|---|---|
| FDmiX | 73.66 ± 0.51 | 0.06 ± 0.04 | -9.10 ± 1.13 | 96.92 ± 0.05 | 252.97 ± 45.88 | 6.40 ± 0.11 |
| DIANT | 76.22 ± 1.48 | 0.06 ± 0.04 | -9.91 ± 1.93 | 96.27 ± 0.18 | 213.64 ± 64.95 | 6.50 ± 0.18 |
| TAMARA | 82.49 ± 1.23 | 0.18 ± 0.02 | -3.75 ± 0.64 | 94.79 ± 0.28 | 156.21 ± 47.53 | 6.35 ± 0.14 |
| Micropore | 87.37 ± 0.11 | 0.04 ± 0.01 | -5.99 ± 2.48 | 93.95 ± 0.06 | 157.27 ± 52.27 | 6.40 ± 0.06 |
| MIVM 3:1 | 99.60 ± 0.99 | 0.25 ± 0.02 | -4.15 ± 0.88 | 93.59 ± 1.52 | 159.33 ± 49.20 | 6.58 ± 0.11 |
| Precigenome | 99.79 ± 1.35 | 0.05 ± 0.02 | -14.54 ± 2.96 | 87.33 ± 1.82 | 104.04 ± 17.07 | 6.25 ± 0.13 |
| Hand mixing | 101.81 ± 1.38 | 0.16 ± 0.02 | -3.36 ± 2.00 | 96.38 ± 0.26 | 100.00 ± 23.75 | 6.57 ± 0.14 |
| Ignite | 102.10 ± 2.64 | 0.22 ± 0.02 | -11.09 ± 2.86 | 91.61 ± 0.16 | 76.28 ± 10.94 | 6.56 ± 0.23 |
| Helix | 161.60 ± 1.01 | 0.16 ± 0.01 | -8.73 ± 0.41 | 63.59 ± 0.99 | 10.72 ± 2.62 | 6.68 ± 0.60 |
| CIJ | 186.50 ± 0.22 | 0.17 ± 0.02 | -8.72 ± 1.59 | 54.58 ± 3.03 | 18.44 ± 4.79 | 6.57 ± 0.04 |
| MIVM 1:1 | 203.10 ± 2.99 | 0.19 ± 0.01 | -12.97 ± 0.78 | 46.49 ± 0.74 | 30.98 ± 26.49 | 6.51 ± 0.19 |

**Figure S 5**. Concentrated FLuc LNP heatmap with standard deviation extending Figure 4

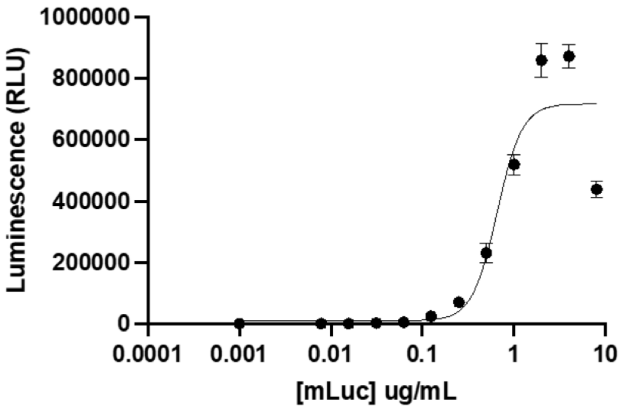


**Figure S 6**. *Dose–response curve of luciferase mRNA LNPs. Luminescence intensity (RLU) was measured across a range of concentrations, and the half-maximal effective concentration ($EC_{50}$) was determined to be 0.65 µg/mL. Data points represent mean ± SD of replicates.*

**Discussion: *in vivo* dosing**

IV-injected LNPs are known to be coated with serum proteins, forming a protein corona that alters their apparent physicochemical properties and governs biodistribution.[1] ApoE adsorption is particularly important, as it directs LNPs to hepatocytes through LDL receptor–mediated uptake, making the liver the primary site of action. Once internalized into cells, LNPs enter the endosomal pathway which gradually acidify during maturation down to pH values of 4–5. This acidification is the key trigger for endosomal escape as the amine groups in ionizable lipids become protonated, which then drives several parallel mechanisms: (1) the accumulation of protons and counterions results in osmotic swelling, (2) the increase in positive charge enhances electrostatic interactions with the negatively charged endosomal membrane, and (3) partial fusion of particle surface with the bilayer lipids.[2] These processes destabilize the endosomal membrane, leading to rupture and release of the encapsulated RNA into the cytoplasm. Once in the cytosol, the higher pH environment weakens the electrostatic interactions between the delivery system and the nucleic acid cargo and facilitating its dissociation from the nanoparticle. The released nucleic acid engages with ribosomes and translated into protein, which resulting in protein (luciferase) expression signaled by luminescence.[3]

**Discussion: TNS assay**

The TNS dye is an anionic fluorescent molecule which fluoresces when it binds to positively charged lipids on an LNP surface. For this assay, LNPs are suspended in different pH conditions. At acidic conditions (low pH), the cationic lipid headgroups become protonated, increasing positive charge on the LNP surface and providing more binding sites for TNS, resulting in higher fluorescent intensity. The pKa value is calculated from the midpoint of the fluorescence drop, which reflects the pH at which half of the ionizable groups are protonated – this can be correlated with the efficiency of LNPs undergoing endosomal escape in cells at acidic conditions, but remaining non-toxic at neutral physiological conditions. The ionizable tertiary amine in ALC-0315 has an intrinsic pKa reported around 6.09.[4] However, when formulated into LNPs with helper lipids and encapsulated RNA, the apparent pKa typically shifts upward, with reported values falling between 6.2 and 6.7, consistent with the optimal window for efficient endosomal escape. Apparent pKa values for each mixer are shown in main body Figure 4c, and curve fits are shown in **Figure S4**.

**Figure S4: TNS assay (apparent pKa) fitting for each mixer, compared to hand mixing**

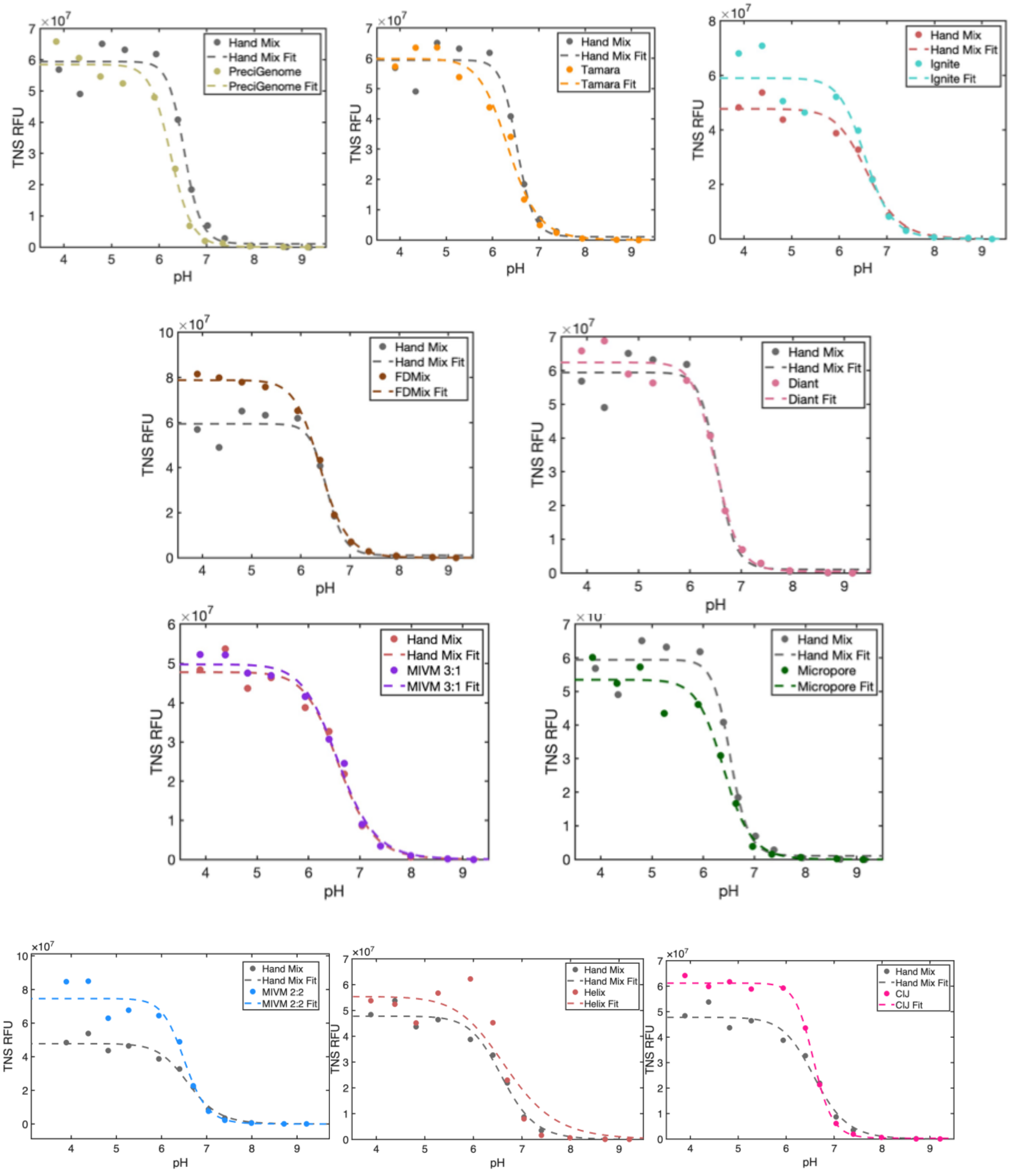


**Figure S 7**. Apparent pKa analysis per mixer compared to hand mixing
